# GlucoCEST MRI for the early evaluation response to chemotherapeutic and metabolic treatments in a murine triple negative breast cancer: a comparison with [^18^F]F-FDG-PET

**DOI:** 10.1101/2021.03.16.432430

**Authors:** Martina Capozza, Annasofia Anemone, Chetan Dhakan, Melania Della Peruta, Martina Bracesco, Sara Zullino, Daisy Villano, Enzo Terreno, Dario Livio Longo, Silvio Aime

## Abstract

**Purpose:** Triple-negative breast cancer (TNBC) patients have usually poor outcome after chemotherapy and early prediction of therapeutic response would be helpful. [^18^F]F-FDG-PET/CT acquisitions are often carried out to monitor variation in metabolic activity associated to response to the therapy, despite moderate accuracy and radiation exposure limit its application. The glucoCEST technique relies on the use of unlabelled D-glucose to assess glucose uptake with conventional MRI scanners and is currently under active investigations at clinical level. This work aims at validating the potential of MRI-glucoCEST in monitoring early therapeutic responses in a TNBC tumor murine model.

**Procedures:** Breast tumor (4T1) bearing mice were treated with doxorubicin or dichloroacetate for one week. PET/CT with [^18^F]F-FDG and MRI-glucoCEST were performed at baseline and after 3 cycles of treatment. Metabolic changes measured with [^18^F]F-FDG-PET and glucoCEST were compared and evaluated with changes in tumor volumes.

**Results:** Doxorubicin treated mice showed a significant decrease in tumor growth when compared to the control group. GlucoCEST imaging provided early metabolic response after three cycles of treatment, conversely, no variations were detect by in [^18^F]F-FDG uptake. Dichloroacetate treated mice did not show any decrease either in tumor volume or in tumor metabolic activity as assessed by both glucoCEST and [^18^F]F-FDG-PET.

**Conclusions:** Early metabolic changes during doxorubicin treatment can be predicted by glucoCEST imaging that appears more sensitive than [^18^F]F-FDG-PET in reporting on early therapeutic response. These findings support the view that glucoCEST may be a sensitive technique for monitoring metabolic response, but future studies are needed to explore the accuracy of this approach in other tumor types and treatments.

## INTRODUCTION

Breast cancer is the most common invasive malignancy in women worldwide, with triple-negative breast cancer (TNBC) accounting for 15% of invasive breast cancers and characterized by high aggressiveness and different sensitivity to chemotherapy [1]. In clinical settings, tumor diagnosis and therapy response are often performed by Positron Emission Tomography (PET) in combination with 2-deoxy-2-^18^F-Fluoro-D-glucose (^18^F-FDG) [2–3] to exploit the elevated glucose uptake in tumors [4]. A change in tumor metabolic activity, as assessed by a decrease of [^18^F]F-FDG uptake, is an important hallmark of treatment efficacy [5–6] and it is associated to pathological responders [7–9].

However, the radioactivity dose and X-ray exposure associated with the [^18^F]F-FDG-PET/CT methodology are source of concerns that limit its use, especially when repeated exams are required [10–11]. Additionally, [^18^F]F-FDG-PET/CT has reasonable sensitivity but low specificity in assessing response to chemotherapy in breast cancer [12].

Therefore, a molecular imaging approach, without the use of ionizing radiation, able to report on the tumor metabolic activity appears most of need. For these reasons, the development of radiation-free MRI methods based on the CEST (Chemical Exchange Saturation Transfer) technique has gained much interest in the biomedical community for investigating tumor metabolism [13–15]. Briefly, a CEST experiment consists in the application of radio frequency pulses at the frequency of the contrast agent protons that are involved in chemical exchange with bulk water protons. Each saturated proton transferred to the bulk water by the chemical exchange is then replaced by an “unsaturated” water proton in a continuous process that, in turn, will result in a detectable decrease of the water resonance [16–18]. MRI glucoCEST is an emerging technique that exploits native glucose for non-invasively tumor characterization both at preclinical [13, 19–22] and clinical levels [23–25]. D-glucose was proposed for the first time in 2012 as CEST contrast agent to discriminate between two breast tumor models with a different tumor metabolism [21]. Although glucose and [^18^F]F-FDG possess a different metabolic fate, with both readily taken up by cancer cells, a good spatial accordance between the [^18^F]F-FDG autoradiography and glucoCEST images was observed in colorectal tumor models [26]. After these first studies, several not metabolizable glucose analogues were proposed as CEST contrast agents for the assessment of tumor metabolism as more closer mimics to the [^18^F]F-FDG [27–31]. However, D-glucose has still a great advantage in terms of safety and bio-tolerability, that pushed the clinical investigations in several cancer patients [23, 32–33], albeit further optimization and some technical challenges need to be solved [34–35]. Several other metabolites, including lactate, can also be detected by this technique or by exploiting dedicated probes [36–39]. Besides tumor detection and staging, the CEST technique can also be applied for therapy response monitoring [39–42]. Recently, glucoCEST imaging was exploited to assess the response to a mTOR inhibitor, rapamycin, in a glioblastoma tumor model [43].

The aim of this study was to investigate whether glucoCEST approach is a valid radiation-free alternative to [^18^F]F-FDG-PET in monitoring the metabolic response to anticancer therapies in a preclinical breast cancer murine model. 4T1 murine cancers were used as a clinical relevant highly metastatic TNBC phenotype and treated with two different therapies, based on the chemotherapeutic doxorubicin or specifically targeting tumor metabolism with dichloroacetate.

Doxorubicin is widely used in the neoadjuvant setting for TNBC as it kills rapidly proliferating cancer cells by inhibiting topoisomerase II [44]. However, [^18^F]F-FDG-PET/CT has a moderate accuracy in predicting the pathological response during chemotherapy in breast cancer patients [45]. Dichloroacetate reverses the Warburg effect in tumor cell metabolism redirecting pyruvate back into the mitochondria and reducing lactate production [46]. Although the effective clinical administration in cancer therapy is still limited to clinical trials, few studies included [^18^F]F-FDG-PET to assess the metabolic response [47].

We evaluated the tumor volume growth in treated and untreated mice and determined whether MRI-glucoCEST or [^18^F]F-FDG-PET are useful for assessing the early treatment response after few cycles of therapy.

## MATERIALS AND METHODS

Materials and methods are presented in the electronic supplementary material (ESM).

## RESULTS

Tumor volume reduction in doxorubicin treated mice compared to control mice was already observed five days after treatment (19 days post-implantation, Fig 1). Differences in tumor volumes between the two groups of mice became more evident and statistically significant after the second doxorubicin dose (292 mm^3^ and 188 mm^3^, P = 0.0004; 430 mm^3^ and 302 mm^3^ P <0.0001, 628 mm^3^ and 321 mm^3^ P <0.0001 for control and treated mice, after 21, 24 and 26 days post tumor implantation, respectively), indicating that the doxorubicin treatment was effective in reducing the tumor growth. Conversely, dichloroacetate was not effective in reducing tumor growth in the investigated murine breast tumor model, with similar tumor volumes between control and treated mice.

**Figure 1.**
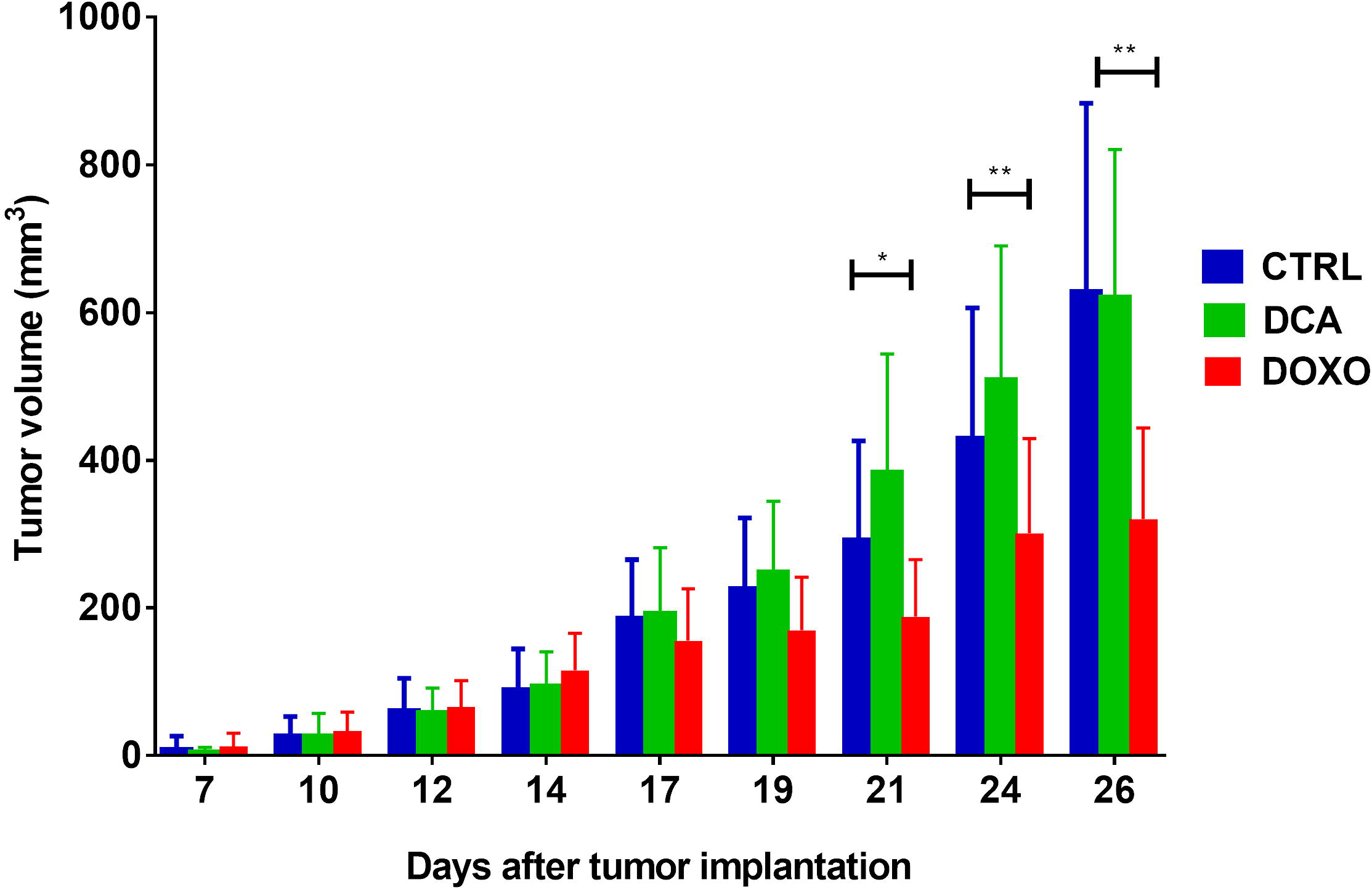
Tumor volume. Bar graph showing tumor volume calculation of 4T1 tumor bearing mice over time for control, doxorubicin and dichloroacetate treated groups (mean ± SD). Two-way ANOVA Bonferroni post, * p=0.0002, ** p<0.0001.

The mean glucoCEST contrast (ΔST%) was similar between pre and post saline treatment in control group (ΔST% = 3.2 ± 1.1 and 2.9 ± 1.2, Fig 2. **(a)**), likewise the fraction pixel that is the number of pixels within the tumor region where glucose was detectable revealed a similar amount of pixels (0.80 ± 0.15 and 0.85 ± 0.16, Fig. 2 **(b)**). There were no significant changes in [^18^F]F-FDG uptake from baseline to post-treatment (%ID/g = 2.0 ± 0.2 and 2.3 ± 0.3, Fig. 3 **(c)**).

**Figure 2.**
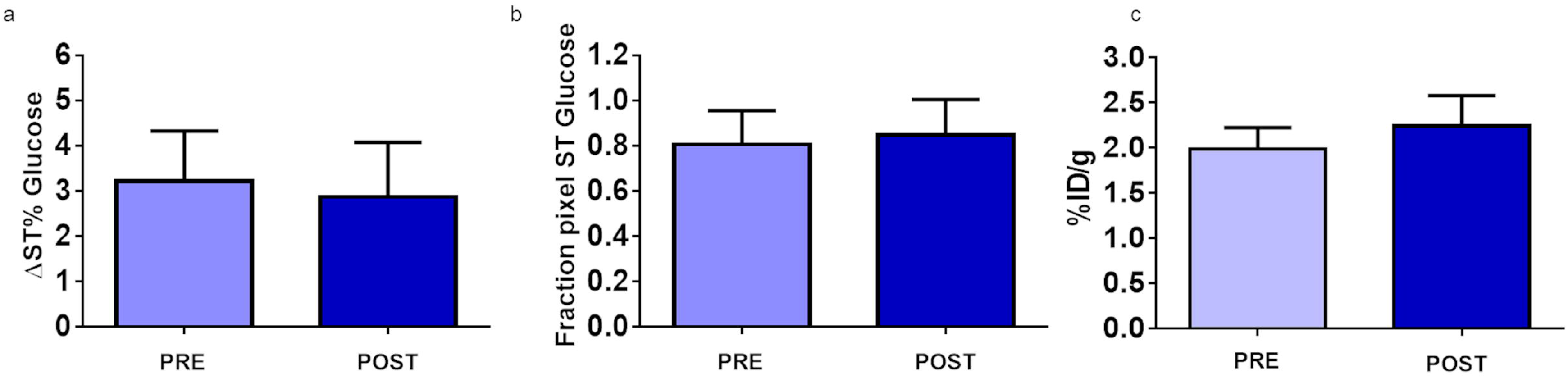
Control group. GlucoCEST and [^18^F]F-FDGsignal measured in the 4T1 tumor model for the control group before (PRE, 14 days after tumor implantation) and after saline treatment (POST, 21 days after tumor implantation). A) Bar graph showing mean ± SD glucoCEST contrast obtained injecting glucose at 3g/Kg dose via intravenous bolus for each 4T1 tumor group. Data are reported as the variation (ΔST%) between the ST effect post-injection minus the ST effect pre-injection. B) Bar graph showing the fraction pixel values (mean ± SD). C) Bar graphs show the average values for the injected dose per gram (%ID/g, mean ± SD) for the ^18^F-FDG-PET study.

**Figure 3:**
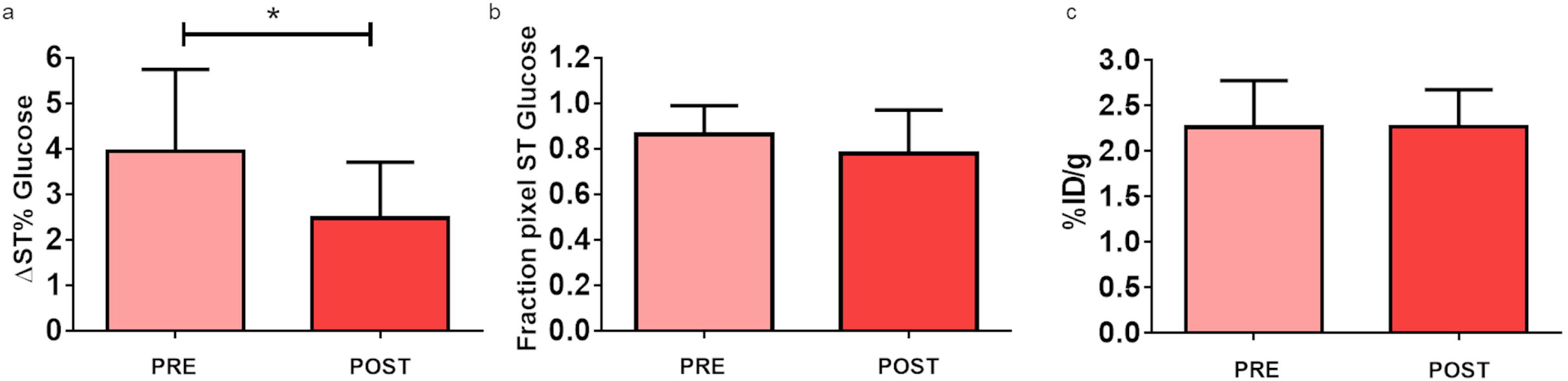
Doxorubicin treated group. GlucoCEST and [^18^F]F-FDGsignal measured in the 4T1 tumor model for the doxorubicin group before (PRE, 14 days post tumor implantation) and after doxorubicin treatment (POST, 21 days post tumor implantation). A) Bar graph showing mean ± SD GlucoCEST contrast obtained injecting glucose at 3g/Kg dose via intravenous bolus for each 4T1 tumor group. Data are reported as the variation (ΔST%) between the ST effect post-injection minus the ST effect pre-injection. Unpaired t-test * p= 0.0334. B) Bar graph showing the fraction pixel values (mean ± SD). C) Bar graphs showing the average values for the injected dose per gram (%ID/g, mean ± SD) for the ^18^F-FDG-PET study

Doxorubicin treated group showed a marked glucoCEST contrast decrease after three doses of doxorubicin (5 mg/Kg ip each). The ΔST% was 4.0% ± 1.8% before treatment and 2.5% ± 1.2% after treatment (p = 0.0334, t-test Fig. 3 **(a)**). The fraction pixel showed a slight but not significant decrease before and after treatment (0.86 ± 0.13 and 0.78 ± 0.19, Fig. 3 **(b)**). Similarly, the [^18^F]F-FDG uptake was constant before and after treatment (%ID/g = 2.3 ± 0.5 and 2.3 ± 0.4, Fig. 3 **(c)**).

The mean glucoCEST contrast detected before and after dichloroacetate treatment was constant (ΔST% = 3.4 ± 1.4 and 3.4 ± 1.5, Fig. 4 **(a)**). The calculated fraction pixel revealed a similar number of pixels where glucose was detectable inside the tumor volume (0.80 ± 0.18 before and 0.89 ± 0.10 after treatment, Fig. 4 **(b)**). Similarly, when comparing the [^18^F]F-FDG uptake values between groups, no differences were observed before and after treatment (%ID/g values were 2.2 ± 0.3 and 2.3 ± 0.5 before and after treatment, Fig. 4 **(c)**). The native D-glucose was well detected in the glucoCEST representative parametric maps (Fig. 6) in control and treated groups at baseline, with a marked decrease in the glucoCEST contrast for the post doxorubicin treated group. Noteworthy, the average signal was lower in doxorubicin treated tumor, but the distribution of the signal was similar between groups as reported by the fraction pixel values. The [^18^F]F-FDG accumulation provided a good visualization of the radiolabelled glucose uptake within the tumor regions, with no appreciable differences after any treatment (Fig. 6). At the end of the therapeutic regimen, the tumor tissue of each group was removed and H&E staining was performed. Fig. S2 in the supplementary (see ESM) shows a similar percentage area of tumor necrosis for all the three groups.

**Figure 4.**
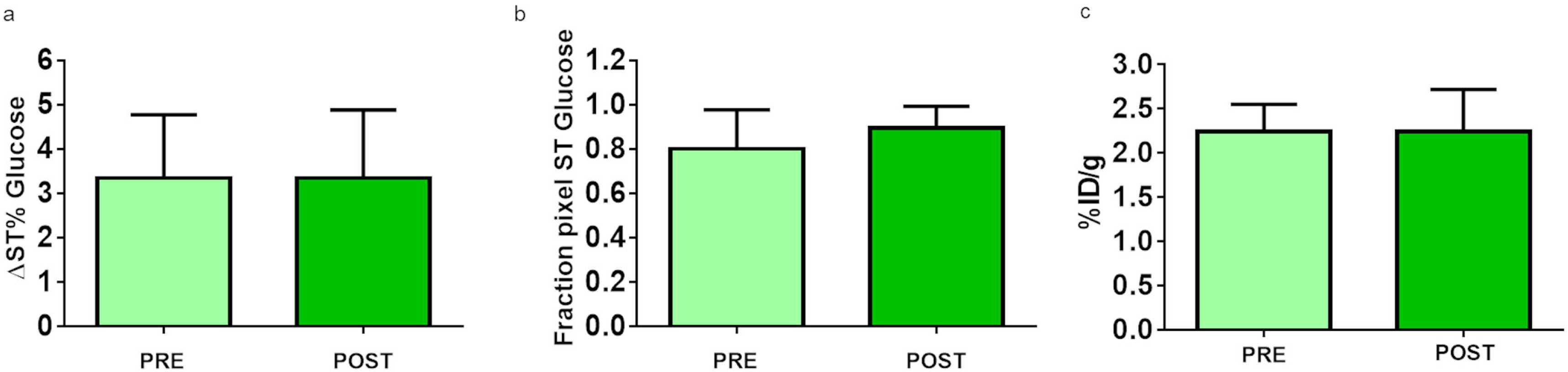
Dichloroacetate group. GlucoCEST and [^18^F]F-FDGsignal measured in the 4T1 tumor model for dichloroacetate group before (PRE, 14 days post tumor implantation) and after dichloroacetate treatment (POST, 21 days post tumor implantation). A) Bar graph showing mean ± SD GlucoCEST contrast obtained injecting glucose at 3g/Kg dose via intravenous bolus for each 4T1 tumor group. Data are reported as the variation (ΔST%) between the ST effect post-injection minus the ST effect pre-injection. B) Bar graph showing the fraction pixel values (mean ± SD). C) Bar graphs showing the average values for the injected dose per gram (%ID/g, mean ± SD) for the ^18^F-FDG-PET study.

**Figure 5.**
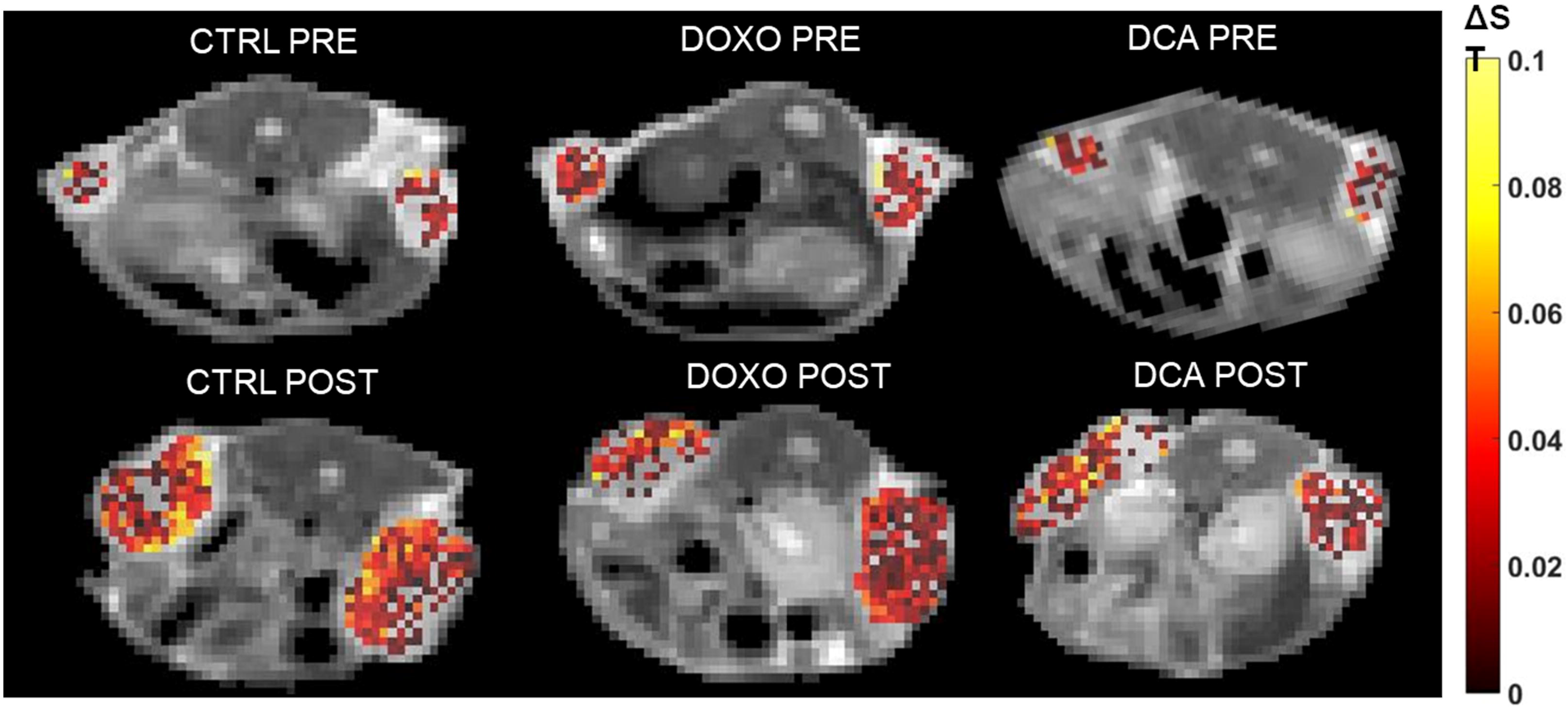
**Representative glucoCEST ΔST% maps** of 4T1 tumor bearing mice for control and treated groups before and after treatment. Data are reported as the difference (ΔST %) between the ST effect before and after the intravenous glucose injection. Parametric maps are overimposed to T2w anatomical images and glucoCEST contrast is shown only in the tumor region.

**Figure 6.**
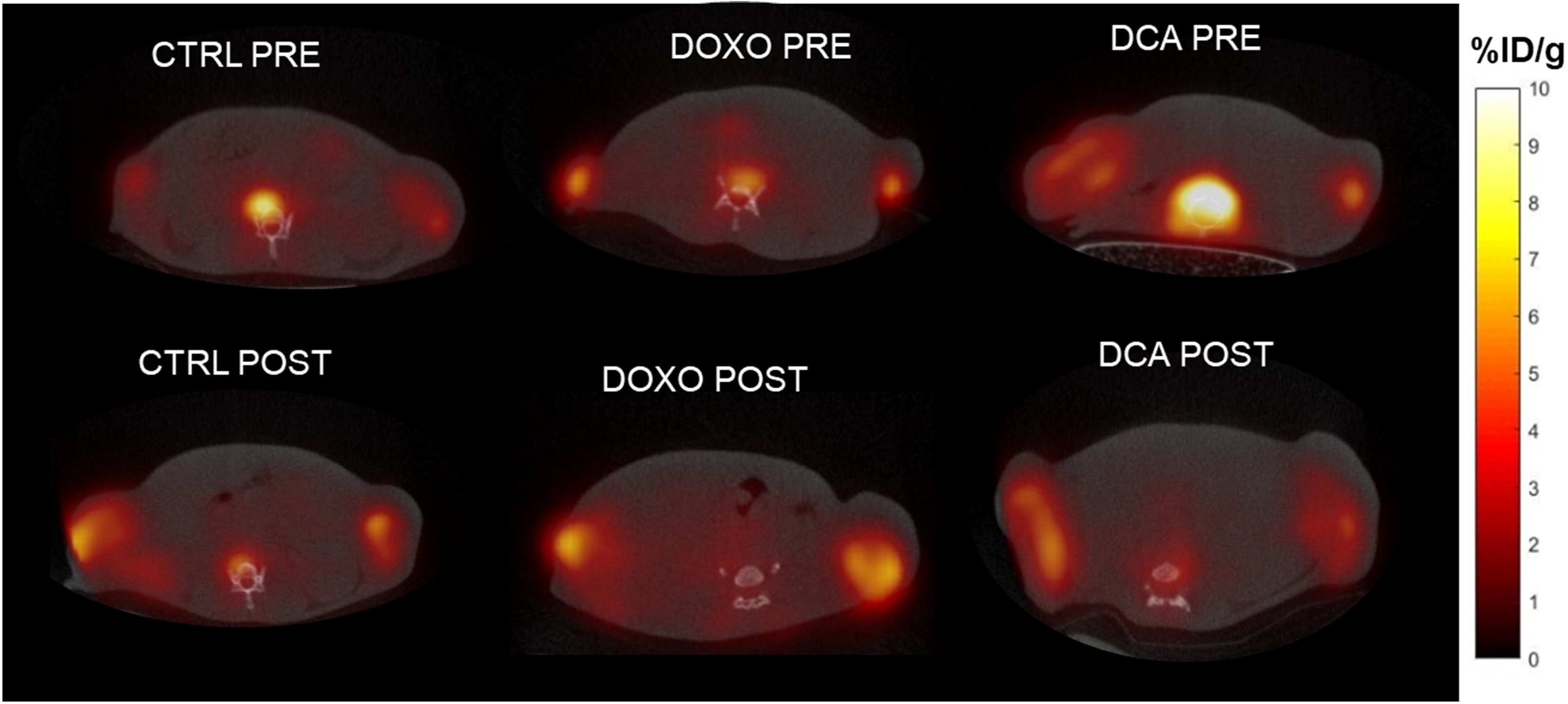
**Representative [^18^F]F-FDG images** of 4T1 tumor bearing mice for control and treated groups before and after treatment. Data are reported as %ID/grams after [^18^F]F-FDG intravenous injection. Fused PET/CT axial view are shown.

## DISCUSSION

The aim of this work was to investigate the capability of the MRI glucoCEST methodology to assess the early therapeutic response to anticancer treatments based on doxorubicin or on dichloroacetate. The glucoCEST results were compared with the [^18^F]F-FDG-PET technique and with the tumor size changes that are the established standard approach for monitoring therapy response in oncology [5–6, 48].

The herein used murine breast (4T1) tumor model resulted responsive to the doxorubicin treatment, with a marked and significant reduction in tumor volume with respect to the control group, while the dichloroacetate treatment was not effective in reducing tumor size. According with the volume measurements, a reduction in the observed glucoCEST contrast for the doxorubicin treated mice when compared to baseline was observed while in dichloroacetate treated mice the glucoCEST contrast did not vary between pre- and post-treatment time points thus in agreement with the absence of tumor size reduction. Conversely, the pixel fraction and the [^18^F]F-FDG-PET approach were not able to report on the effects associated with both doxorubicin and dichloroacetate treatments.

Doxorubicin is a potent anthracycline antibiotic and DNA intercalator that inhibiting topoisomerase II slows down cancer cell growth[49–50]. Doxorubicin is responsible for cell apoptosis, autophagy, senescence and necrosis[51] and it is effective in the treatment of leukaemia and Hodgkin’s lymphoma [52–53], as well as several other cancers [54–55]. Our results reporting the poor diagnostic potential of [^18^F]F-FDG-PET are in agreement with observations performed in different cancer models where doxorubicin inhibited tumor growth but the [^18^F]F-FDG-PET parameters were not significantly affected[56–59]. As hypothesized by Malinen et al[57], in a complex environment such as the tumor, different factors could affect the [^18^F]F-FDG uptake. For example, decreases of tumor metabolism and increases of vascularization may balance out and results in a constant [^18^F]F-FDG uptake. In another study, murine mammary tumors were treated with doxorubicin and, in spite of a decrease of [^18^F]F-FDG uptake after 1 day of treatment, a transient increase was reported 7 days later[60]. The limits of [^18^F]F-FDG-PET in assessing treatment response in comparison to novel imaging approaches based on tumor acidosis have been reported also following metformin treatment[61]. These results suggest the need of reconsidering the accuracy of [^18^F]F-FDG-PET as biomarker of doxorubicin treatment efficacy in clinical routine [62–63] to avoid the risk of overestimate or underestimate the therapeutic response.

Dichloroacetate shifts metabolism from glycolysis to oxidative phosphorylation, hence reducing lactate production[64–65]. Several clinical trials have already been performed to test dichloroacetate activity as an anticancer agent[66]. However, at preclinical level, the effects of dichloroacetate appear to be dependent on the investigated cancer cell line and imaging technology [67]. A decrease of [^18^F]F-FDG uptake was observed in patients with solid tumors treated with dichloroacetate [47] while in human breast cancer (MDA-MB-231) and human cervical cancer (SiHa) mouse models the [^18^F]F-FDG uptake is not affected by dichloroacetate treatment[68]. In a glioma model the early metabolic response to dichloroacetate treatments has been monitored with hyperpolarized ^13^C-pyruvate MRI showing marked metabolic changes in lactate and bicarbonate levels [69]. Dichloroacetate in a TS/A breast cancer xenograft model had no overall effect in tumor size but a marked tumor pH increase was observed in treated tumors by MRI-CEST pH imaging that correlated with a lactate decrease at earlier time points[70]. In a U87MG glioma model the intracellular acidification was detected even one hour post dichloroacetate administration using MRI-CEST[71]. Our results showed that the dichloroacetate treatment was not effective in the 4T1 tumor model, since neither tumor volume, nor ^18^F-FDG-PET uptake and glucoCEST uptake were affected.

The fraction pixel metric, an indirect measurement of tumor vascularization based on the contrast agent detectability, highlighted a similar vascularization between control and treated groups. Also the histological evaluation showed a comparable extension of the necrotic areas among the three groups. We can ascribe the larger necrosis in the control and dichloroacetate groups due to the larger tumor volumes, in contrast to the smaller tumor volumes for the doxorubicin group. Even though it has been reported that doxorubicin treatment induces necrosis[51], several studies on the 4T1 breast cancer model showed a similar necrotic area between treated and control tumors[72–73].

GlucoCEST imaging suffers of low contrast enhancement, hence requiring high magnetic fields to improve the detectability and the sensitivity of this approach [33–34, 74]. However, a great deal of effort is putting into sequences and irradiation pulse optimization in order to improve glucoCEST detection[75–76].

## CONCLUSION

To the best of our knowledge, this is the first preclinical evaluation of the ability of glucoCEST imaging for assessing the early treatment efficacy in two different therapies with a direct comparison with [^18^F]F-FDG-PET technique. Overall, the findings reported herein support the use of glucoCEST imaging as a valid alternative to [^18^F]F-FDG-PET for assessing early treatment response to conventional chemotherapy. However, additional preclinical studies are needed to further determine the capability of glucoCEST imaging in assessing therapeutic response in other tumor models and therapeutic regimens.

## ACKNOWLEDGEMENTS

We gratefully acknowledge support from the European Union’s Horizon 2020 research and innovation program (Grant Agreement No. 667510, GLINT project) to SA, ET, MC, AA, SZ, DV. MC was supported by fellowship from the Italian Association for Cancer Research (AIRC ID 24104). The Italian Ministry for Education and Research (MIUR) is gratefully acknowledged for yearly FOE funding to the Euro-BioImaging Multi-Modal Molecular Imaging Italian Node (MMMI).

## AUTHOR CONTRIBUTION

Conceptualization, S.A., D.L.L. and E.T.; design of the work, D.L.L., M.C., A.A.; data acquisition M.B.; M.D.P. M.C.; acquisition analysis M.B., C.D., D.V., S.Z.; writing—original draft, M.C., A.A,; writing—review and editing, D.L.L., E.T., S.A.

