## supplementary for "GlucoCEST MRI for the early evaluation response to chemotherapeutic and metabolic treatments in a murine triple negative breast cancer: a comparison with [^18^F]F-FDG-PET"

Dario Livio Longo, Istituto di Biostrutture e Bioimmagini (IBB), Consiglio Nazionale delle Ricerche (CNR), Via Nizza 52, 10126, Torino, Italy

**Chemicals**

Glucose solution for intravenous administration was prepared by dissolving D-glucose (Sigma-Aldrich, St. Louis, MO, USA) in saline solution to obtain a 3 M solution. Before the use, the solution was filtered with a 200 nm membrane filter in order to preserve the suspension from bacterial contamination.

[$^{18}$F]F-FDG (GLUSCAN® 600 MBq/mL) was purchased by Advanced Accelerator Applications, a Novartis Company (Saint-Genis-Pouilly, France). The Doxorubicin solution for intraperitoneal injection was prepared by dissolving Doxorubicin hydrochloride powder (European Pharmacopoeia (EP) Reference Standard, Sigma-Aldrich, St. Louis, MO, USA) in saline solution to obtain a concentration of 1 mg/ml. In order to avoid precipitation, the powder was dissolved right before administration.

Dichloroacetate solution for intraperitoneal administration was prepared by dissolving sodium dichloroacetate powder (Sigma-Aldrich, St. Louis, MO, USA) in saline solution to reach the concentration of 50 mg/mL, whereas for the oral administration, the powder was dissolved in water to reach a concentration of 0.45g/L.

**Cell line**

4T1 cells were obtained from American Type Culture Collection (ATCC, Manassas, VA, USA). Cells were grown in RPMI 1640 (Euroclone S.p.A, Pero, Italy) medium supplemented with 10% FBS (Euroclone S.p.A), 100U/mL Pen/Strep (Euroclone S.p.A) and 2 mM L-Glutamine (Euroclone S.p.A). The cells were grown at 37°C in a humidified atmosphere containing 5% $CO_2$.

**Tumor model**

Female BALB/c mice were purchased by Envigo RMS, S.r.l., Udine Italia. All the procedures involving the animals were carried out according to the national and international laws on experimental animals (L.D. 26/2014; Directive 2010/63/EU). Tumor model was obtained by subcutaneous injection of 4.0 x $10^4$ 4T1 cells into both mice flanks (n=42) under sevoflurane (Alcyon Italia S.P.A., Cherasco, Italy) anesthesia.  Length and width of the tumors were measured using a caliper three times a week starting from baseline up to 26 days and the volumes were calculated using the equation ($Width^2 \times Length$)/2.

**Experimental design**

**Figure S1** shows a graphical representation of the experimental setup. Fourteen days after inoculation, all the mice included in the study underwent glucoCEST MRI and [$^{18}$F]F-FDG-PET examinations before the treatment (PRE). Then, mice were randomly divided into three groups. Control group (n=13) was intraperitoneally injected with a vehicle solution at the same time points of the treated groups. Doxorubicin group received 4 doses of 5mg/Kg bw of doxorubicin (n=17) through intraperitoneal injection with an interval of 2-3 days (15, 18, 20 and 22 days post tumor implantation). A third group of mice was treated with dichloroacetate (n=12) via intraperitoneal route at a dose of 200 mg/kg/day and orally with mice having access to a 0.45g/ml solution ad libitum for 12 days(Anemone et al. 2017). One week after the starting of the treatment (i.e. 21 days post-implantation), post-treatment MRI and PET examinations were performed. GlucoCEST images were acquired for 6 mice for both control and doxorubicin-treated group and for 5 mice in the dichloroacetate-treated group. $^{18}$F-FDG-PET/CT images were acquired on a separate group of mice (n=6 for each group) before and after treatment at the same time points.

**GlucoCEST MRI imaging protocol and analysis**

MR images were acquired using a 7 T Bruker Pharmascan (Bruker Biospin, Ettlingen, Germany) scanner equipped with a 30 mm 1H quadrature coil. Before imaging, mice were anesthetized by isoflurane (Alcyon Italia S.P.A., Cherasco, Italy) vaporized with $O_2$. Isoflurane was used at 3.0 % for induction and at 1.0 % to 2.0 % for maintenance, placed on the MRI bed and an air pillow placed underneath the animal (SA Instruments, Stony Brook, NY; USA) monitored breath rate. The tail vein was cannulated with a catheter to administer glucose at a dose of 3 g/kg bw through a 27-gauge needle. After the scout image acquisition, $T_{2w}$ anatomical images were acquired with a RARE sequence (TR = 4 seconds, TE = 3.7 milliseconds, NA = 1, slice thickness = 1.5 mm, FOV = 30 × 30 mm, matrix size = 256 × 256). The same geometry was used for the following glucoCEST acquisitions by irradiating with a single, continuous pre-saturation block pulse of 2 μT applied for 5 sec. The saturation frequency offset ranged from -10 to 10 ppm with a frequency resolution of 0.2 ppm.

In-house MATLAB (The Mathworks, Inc., Natick, MA, USA) scripts have been used to process all the CEST images. Anatomical and Z-spectrum images were first segmented by using an intensity-threshold filter(Chu and Hamarneh 2006). The Z-spectra were interpolated, on a voxel-by-voxel basis, by smoothing splines(Stancanello et al. 2008) to identify the correct position of the bulk water, thus removing artifacts arising from $B_0$ inhomogeneity. On this basis, the interpolated Z-spectrum was shifted so that the bulk water resonance corresponds to the zero frequency and corrected intravoxel saturation transfer (ST) effects were calculated(Terreno et al. 2009). Then, a second filter was applied to remove CEST effect arising from noisy data, calculating the coefficient of determination $R^2$ for the interpolating curve to take into account the signal-to-noise ratio of single voxels (noisy Z-spectra present low $R^2$ values). Only voxels with high $R^2$ (>0.99) were considered in the ST calculation. The ST effect for glucose was estimated from the expression:

$$ST = \frac{S(-1.2ppm) - S(1.2ppm)}{S_0}$$

where $S_0$ was the signal at -10 ppm.

Results are reported as:

$$\Delta ST\% = (\text{ST post-injection} - \text{ST pre-injection}) * 100$$

The fraction pixel reports on the percentage of pixels showing a positive ΔST% in the manually-defined tumor region of interest (ROI) in MATLAB.

**[$^{18}$F]F-FDG-PET imaging protocol and analysis**

Animals were fasted overnight before the intravenous [$^{18}$F]F-FDG injection (5.48 ± 0.19 MBq/mouse). The injected dose was calculated by the difference of the radioactivity in the syringe before and after the administration, as measured by a dose calibrator (IsoMED 2010, MED Nuklear-Medizintechnik Dresden GmbH). Animals were anesthetized with isoflurane (3 % for induction, 1 %

- 1.5 % for maintenance, in oxygen) and then placed in a dedicated small-animal multimodality scanner (Triumph II, TriFoil Imaging, Chatsworth, CA), equipped with inhalation anesthesia and heating pad. Instrument calibration was performed with phantoms containing small known amounts of radioactivity. A whole-body CT scan was performed immediately before the PET acquisition in order to provide anatomical information (512 projections, 1 frame per projection, 75 kV peak tube voltage, 150 mA tube current, 230 ms exposure time). PET static acquisition was performed for 30 minutes starting 45 minutes after [$^{18}$F]F-FDG administration. PET and CT images were reconstructed using the LabPET software (TriFoil Imaging, Chatsworth, CA). All images were corrected for decay and tissue attenuation, and the CT data were used to provide attenuation correction. The PET data were reconstructed in a single frame using a 3D algorithm (Ordered Subset Expectation Maximization, OSEM-3D) with 8 subsets and 10 iterations. Reconstructed and co-registered PET/CT images were quantitatively evaluated using the AMIDE software package (http://amide.sourceforge.net/)(Loening and Gambhir 2003), by drawing volumes of interests (VOIs, thickness 1 mm) over tumors on axial images. PET data are expressed as percentage of injected dose per gram (%ID/g).

**Histological analysis**

Two mice per group (n=4 tumors/group) were euthanized immediately after the last imaging session, 21 days after the tumor implantation, for histological analysis.  After euthanasia tumors were collected and fixed in a 10% formalin solution. Tissues that have been formalin-fixed and paraffin-embedded were sectioned (5 μm thickness section) and staining procedure with hematoxylin and eosin (H&E) was used. The necrotic area of the histological sections was imaged by light microscopy (1.25×). The percentage of necrosis per total area was determined using ImageJ software.

**Statistical analysis**

Data are expressed as mean ± SD in all cases. The statistical significance of results was determined by a 2-way ANOVA with Bonferroni post-hoc testing for tumor volume measurements, while for glucoCEST and [$^{18}$F]F-FDG values an unpaired t-test was applied. Statistical analysis was performed with Graphpad Prism version 7.00 (La Jolla, California, USA).

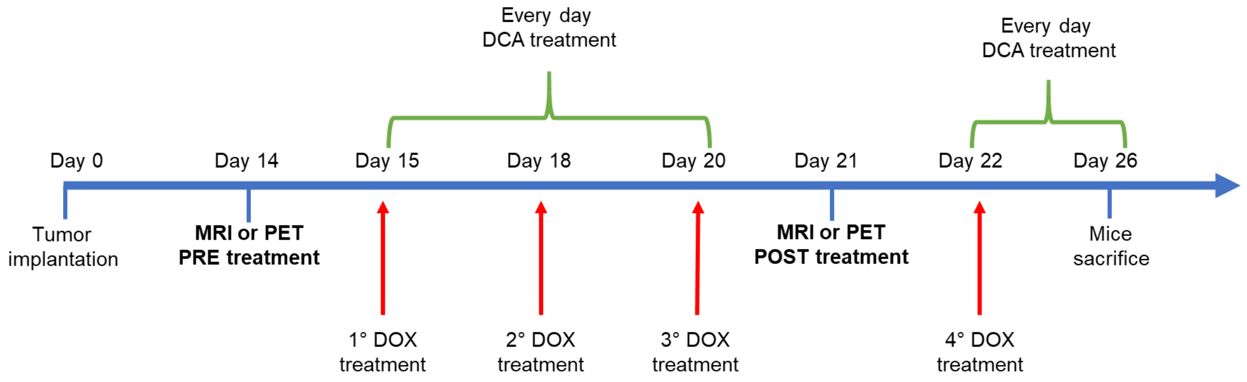


**Figure S1. Experimental design.** 14-days after inoculation, mice underwent imaging acquisition. Mice were randomly divided into three groups. Control, Doxorubicin (4 doses of 5mg/Kg b.w. i.p.) and Dichloroacetate (200 mg/kg/day i.p. and 0.45g/ml solution ad libitum). One week after the starting of the treatment (i.e. 21 days post-implantation), post-treatment imaging examinations were performed.

.

**Figure S2**

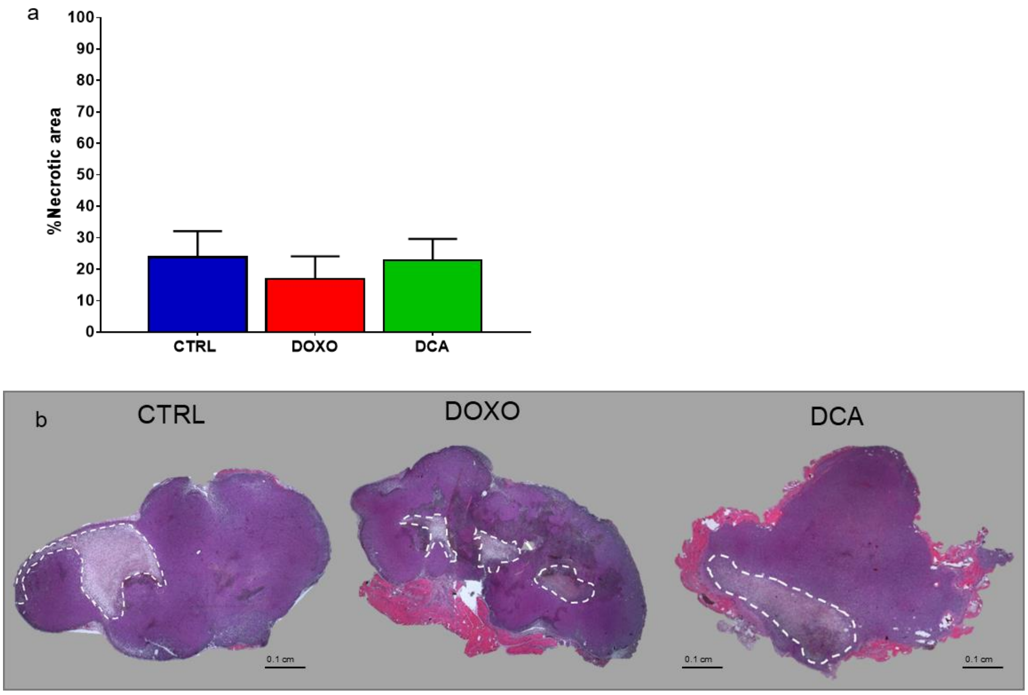


**Figure S2. Representative H&E images** of 4T1 tumor bearing mice for control and treated groups after end of the treatment. a) Quantification of necrotic areas in 4T1 tumor sections (n=4). b) Whole mount sections of 4T1 tumors display a similar percentage of necrosis after treatment (dotted line highlights the necrotic area).
